# Topological constraints in early multicellularity favor reproductive division of labor

**DOI:** 10.1101/842849

**Authors:** David Yanni, Shane Jacobeen, Pedro Márquez-Zacarías, Joshua Weitz, William C. Ratcliff, Peter J. Yunker

## Abstract

Reproductive division of labor (e.g., germ-soma specialization) is a hallmark of the evolution of multicellularity, signifying the emergence of a new type of individual and facilitating the evolution of increased organismal complexity. A large body of work from evolutionary biology, economics, and ecology has shown that specialization is beneficial when further division of labor produces an accelerating increase in absolute productivity (i.e., productivity is a convex function of specialization). Here we show that reproductive specialization is qualitatively different from classical models of resource sharing, and can evolve even when the benefits of specialization are saturating (i.e., productivity is a concave function of specialization). Through analytical theory and evolutionary individual based simulations, our work demonstrates that reproductive specialization is strongly favored in sparse networks of cellular interactions, such as trees and filaments, that reflect the morphology of early, simple multicellular organisms, highlighting the importance of restricted social interactions in the evolution of reproductive specialization. More broadly, we find that specialization is strongly favored, despite saturating returns on investment, in a wide range of scenarios in which sharing is asymmetric.

**Significance Statement:** During the evolution of multicellularity, previously independent cells integrate to form a new organism. A critical step in this major evolutionary transition is the origin of reproductive specialization (e.g., germ-soma differentiation). It is widely thought that complete specialization, like many other types of trade, will only evolve when the payoff from differentiation accelerates with further investment. Here we show that reproductive specialization is a special case of asymmetric trade that is adaptive under a far broader set of conditions, and is particularly promoted in multicellular organisms growing with tree-like topologies. This morphology class readily evolves in nascent multicellular organisms, and appears to be ancestral in most lineages that have evolved complex multicellularity, suggesting a causal role between topology and emergent complexity.

---

**T**he evolution of multicellularity set the stage for unprecedented increases in organismal complexity (1, 2). A key factor in the remarkable success of multicellular strategies is the ability to take advantage of within-organism specialization through cellular differentiation (3–5). Reproductive specialization, which includes both the creation of a specialized germ line during ontogeny (as in animals and volvocine green algae) and functional differentiation into reproductive and non-reproductive tissues (as in plants, green and red macroalgae, and fungi), may be especially important (6–11). Reproductive specialization is an unambiguous indication that biological individuality rests firmly at the level of the multicellular organism (12, 13), and is thought to play an important role in spurring the evolution of further complexity by inhibiting within-organism (cell-level) evolution (14) and limiting reversion to unicellularity (15). Despite the central importance of reproductive specialization, its origin and further evolution during the transition to multicellularity remain poorly understood (16).

The origin of specialization has long been of interest to evolutionary biologists, ecologists, and economists. A large body of theory from these fields shows that specialization pays off only when it increases total productivity, compared to the case where each individual simply produces what they need (1, 17–26). Certain types of trading arrangements maximize the benefits of specialization; highly reciprocal interactions, which facilitate exchange between complementary specialists, amplify cooperation (27, 28). Still, previous work finds that even when groups grow in an ideal spatial arrangement, increased specialization and trade is only favored by natural selection when productivity increases as an accelerating function of the degree of specialization, i.e., productivity is a convex, or superlinear, function of the degree of specialization. Conversely, saturating functional returns (i.e., productivity is a concave, or sub-linear, function of the degree of specialization) should inhibit the evolution of specialization (6–11).

Reproductive specialization differs from classical models of trade in several key respects. Trade between germ (reproductive) and somatic (non-reproductive) cells is intrinsically asymmetric, because the cooperative action, multicellular replication, is not a product that is shared evenly. Selection acts primarily on the fitness of the multicellular group as a whole, which need not be the same as the mean fitness of its component cells (29). As a result, optimal specialization can result in behaviors that reduce the short-term fitness of some cells within the multicellular group (7, 10), often manifest as reproductive altruism.

Understanding the evolution of cell-cell trade, a classic form of social evolution (30), requires understanding the extent of between-cell interactions. Network theory has proven to be an exceptionally powerful and versatile technique for analyzing social dynamics (31, 32), and indeed, is uniquely well suited to understanding the evolution of early multicellular organisms. When cells adhere through permanent bonds, sparse network-like bodies (i.e., filaments and trees) often result. This mode of group formation is not only common today among simple multicellular organisms ((33, 34)), but is the dominant mode of group formation in the lineages evolving complex multicellularity (i.e., plants, red algae, brown algae, and fungi, but not animals).

In this paper, we develop and investigate a model for how the network topology of early multicellular organisms affects the evolution of reproductive specialization. We find that under a broad class of sparse networks, complete functional specialization can be adaptive even when returns from dividing labor are saturating (i.e., concave / sub-linear). Sparse networks impose constraints on who can share with whom, which counterintuitively increases the benefit of specialization (16). By dividing labor, multicellular groups can capitalize on high between-cell variance in fitness, ultimately increasing group-level productivity. Further, we consider group morphologies that naturally arise from simple biophysical mechanisms and show that these morphologies strongly promote reproductive specialization. Our results show that reproductive specialization can evolve under a far broader set of conditions than previously thought, lowering a key barrier to major evolutionary transitions.

## Model

Reproductive specialization can be modeled as the separation of two key fitness parameters, those related to either viability or fecundity, into separate cells within the multicellular organism (13, 35). Thus, we consider a model of clonal cells living within groups that each invest resources into viability and fecundity. We let *v* denote each individual’s investment into viability, and *b* denote each individual’s investment into fecundity. Each individual’s total investment is constrained so that *v* + *b* = 1. However, an individual’s return on its investment is in general nonlinear. Here, we let *α* represent the ‘return on investment exponent’: by tuning *α* above and below 1.0, we can simulate conditions with accelerating and saturating (i.e., convex and concave, or super- and sub-linear) returns on investment, respectively. We let 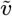 and 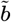 represent a cell’s return on viability and fecundity investments, respectively. Following Michod (35, 36), we calculate an individual cell’s reproductive output as a multiplicative function of 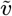 and 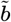 (thus, both functions must be positive for an individual to have nonzero fitness). A single cell’s reproduction rate is 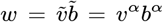. At the group level, fitness is the total contribution of all cells in the group toward the production of new groups (i.e. group level reproduction). The group level fitness is then the sum of 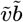 over all cells).

Finally, cells may share the products of their investment in viability with other cells to whom they are connected. For instance, filamentous fungi form networks with cytoplasmic continuity allowing efficient nutrient translocation between cells (37, 38). For a given group, the details about who may share with whom, and how much, is encoded in a weighted adjacency matrix **c**. The element *c*_*ij*_ defines what proportion of viability returns individual *i* shares with individual *j*. Each individual counts itself among its neighbors and always shares a positive portion of viability returns with itself, so that *c*_*ii*_ > 0. Furthermore, since an individual cannot share more viability returns than the total it possesses, we have 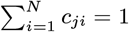 for a group of *N* cells. The total proportion of viabi1lity returns that cell *j* shares with all neighbors is 1. For the networks we consider, each individual takes a fraction *β* of its viability returns and shares that fraction equally among all of its *n*_*i*_ neighbors (including itself), and keeps the rest of its returns 1 − *β* for itself. Therefore individual *i* keeps a total fraction of 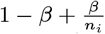 of its returns for itself and gives 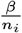 to each of its non-self neighbors. This means the total amount of returns kept by individual *i* depends on *both* the network topology and *β*. When *β* = 0 there is no sharing, and when *β* = 1 individuals share everything equally among all connections. We refer to *β* as interaction strength. A given group topology (unweighted adjacency matrix) and *β* completely specify **c**.

Within a group of *N* cells, the overall returns on viability that a given cell enjoys, then, comprises its own returns as well as whatever is shared with it by other members of the group. This can be written as 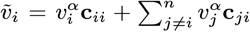, or equivalently, 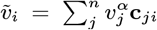. Note that this is a column sum, since it describes the total *incoming* viability returns an
individual receives as a result of toil and trade. Therefore, we write the group level reproduction rate (i.e., the group fitness) for a group of *N* individuals as

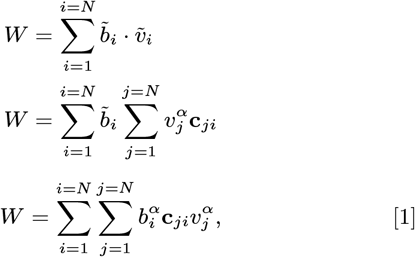

where all three of the above equations are equivalent. We investigate evolutionary outcomes under this definition of group level fitness for groups with different topologies (who shares with whom), and in scenarios with various return on investment exponents *α*.

## Results

### Fixed resource sharing

We first consider cases wherein cells within a group share across fixed intercellular interactions. Individual *i* shares 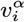 equally among interaction and self terms. In each case we vary the return on investment exponent, *α*, between 0.5 and 1.5, and the interaction strength, *β*, between 0.0 and 1.0, both in increments of 0.1. For each combination of topology, *α*, and *β*, the group investment strategy (*v*_*i*_ for all *i*) was allowed to evolve for 1000 generations.

We begin with simple topologies: groups with no connections and groups that are maximally connected. They represent, respectively, the case in which all individuals within the group are autonomous and the case in which every individual interacts with all others (i.e. a ‘well-mixed’ group). In the absence of interactions, individuals cannot benefit from functions performed by others and therefore must perform both functions *v* and *b*; hence specialization is not favored, and does not evolve. In the fully connected case, a high degree of specialization is observed for many values of *α* and *β* (Figure 1a). Consistent with classic results (6–11), specialization is only achieved in the fully connected case for *α* > 1.

**Fig. 1.**
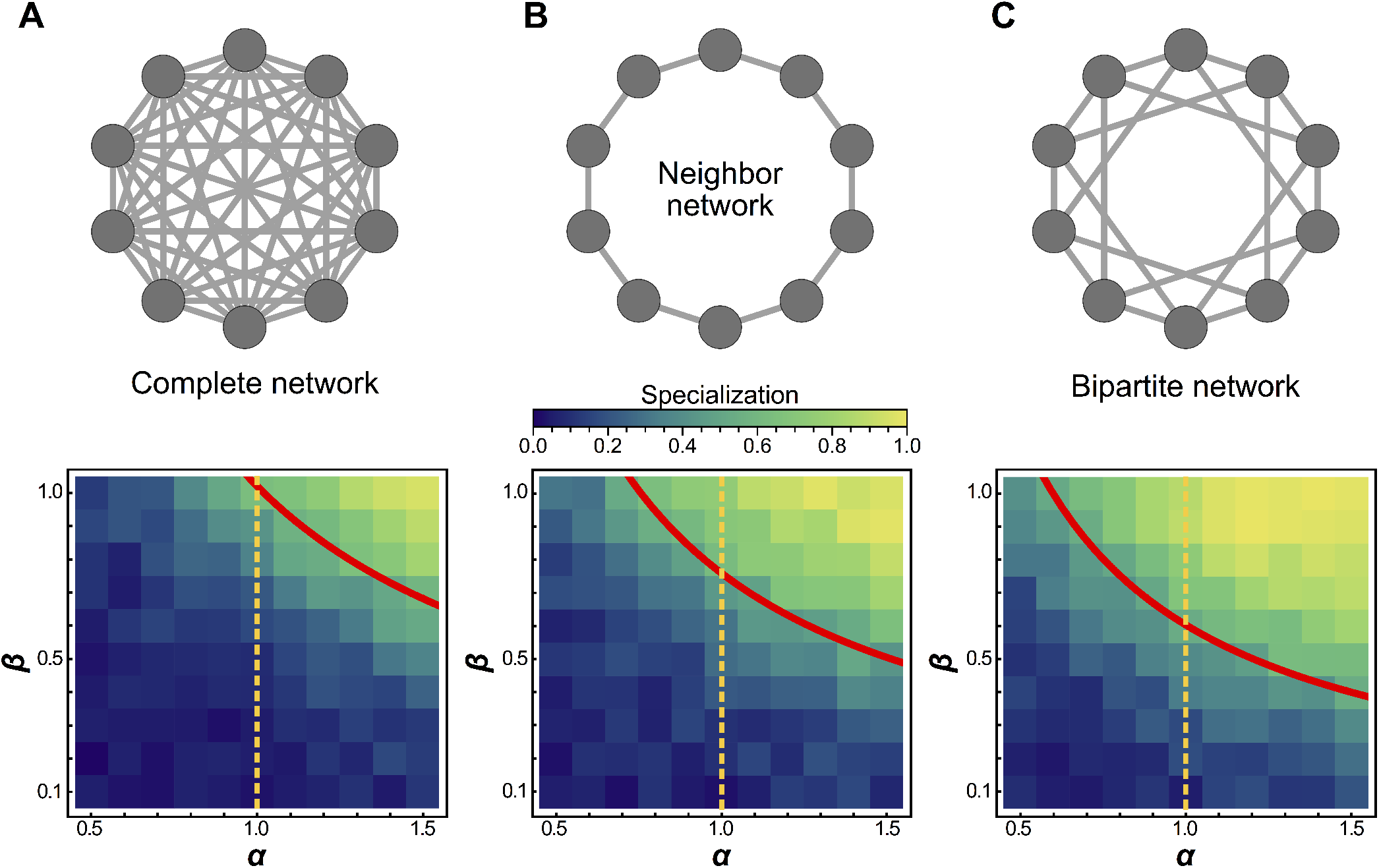
Schematic of topology for a simplified six individual group (first row), and mean specialization as a function of specialization power *α* and interaction strength *β* across the entire population. (a) When each individual in the group is connected to all others, specialization is favored only when *α >* 1. (b) For the nearest neighbor topology, specialization is favorable for a wider range of parameters, including for some values of *α <* 1. Specifically, specialization is advantageous when 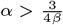 (see Table 1 (c) Connecting alternating specialists creates a bipartite graph which maximizes the benefits of specialization and the range of parameters for which it is advantageous. In this case specialization is favorable wherever 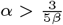. The red curves represent analytical predictions for *α*\*, the lowest value of *α* for which complete generalization is disfavored, and the orange vertical lines are at *α* = 1 to guide the eye. While analysis shows that *some* degree of specialization must occur in the regime upward and to the right of the red curves, simulations reveal that when complete generalization is disfavored complete specialization *is* favored in these networks.

Next, we consider a simple sparse network in which each individual within a group is connected to only two other individuals, forming a complete ring (Figure 1b); we refer to this as the neighbor network. Surprisingly, preventing trade between most individuals encourages division of labor. We find that specialization evolves even when *α <* 1.0, i.e., when the returns on investment are saturating or concave. In our simulations, this topology leads to alternating specialists in viability and fecundity (Figure 1b). Analytically, we find that this topology always favors at least some degree of specialization whenever 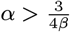.

We next study a network with individuals that can be separated into two disjoint sub-groups, where every edge of the network connects an individual in one sub-group to an individual in the other sub-group and no within sub-group connections exist, i.e., a bipartite graph (Figure 1c).

We can analytically determine under what conditions complete generalization is optimal. The complete generalist investment strategy is where every individual in the group invests equally into viability and fecundity, defined as: 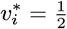. for all *i* For these simple topologies the complete generalist strategy is either a maximum or a saddle point, depending on the values of *α* and *β*. Complete generalization is only favored when the Hessian evaluated at the generalist investment strategy 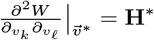 is negative definite, i.e. all of its eigenvalues are negative. Table 1 shows the largest eigenvalues of the Hessian for these topologies. When *α* and *β* are chosen so that the largest eigenvalue becomes non-negative, complete generalization cannot maximize fitness.

**Table 1.**
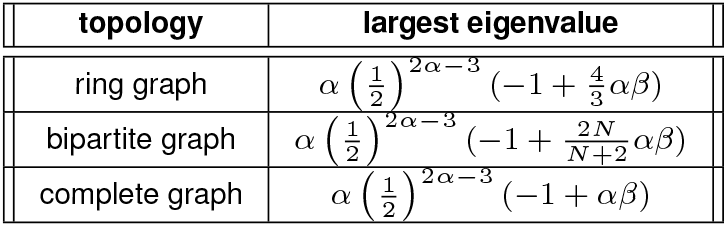
Largest eigenvalue of the Hessian evaluated at the generalist critical point as a function of *α,β*, and *N* for three topologies.

**Table 2.**
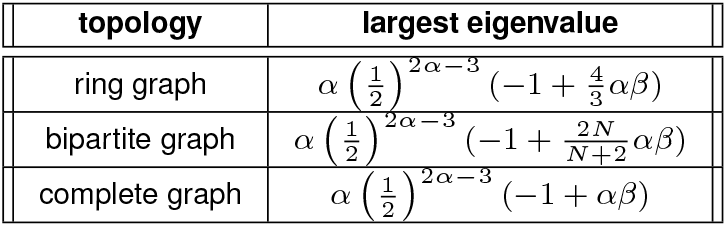
Largest eigenvalue of the Hessian evaluated at the generalist critical point as a function of *α*, *β*, and *N* for three topologies. When the group size *N* = 4, the bipartite graph coincides with the ring graph, and indeed the eigenvalues agree. Similarly, when *N* = 2 the bipartite graph coincides with the complete graph and the eigenvalues agree. The interesting domain of *αβ* is (0, 1], so for the complete graph H* is always negative definite. However, the bipartite and ring graphs show regions where the generalist strategy is *not* stable.

While we have not analytically shown where the fitness maximum occurs in cases where the generalist strategy becomes a saddle point, evolutionary simulations (Figure 1) suggest that when complete generalization is not a fitness maximum, a high degree of (or even complete) specialization typically *does* maximize fitness.

In all cases in which complete specialization is achieved in evolutionary simulations, the self-fitness terms of the viability specialists go to zero, as they cannot reproduce on their own. Furthermore, the fecundity specialists are entirely reliant on the viability specialists for their survival; if viability sharing were suddenly prevented, their fitness would also be zero. This amounts to complete reproductive specialization (6, 30, 35).

### Evolving resource sharing

Until now, sharing has been included in every intercellular interaction within groups. Here, we consider the case in which there is initially no sharing, and sharing must evolve along with specialization. These simulations begin with no resource sharing (i.e., *β* = 0); during every round, each group in the population has a 2% chance that one of its cells will mutate and *β* will change. The new *β* value is chosen from a truncated Gaussian with standard deviation of 10% of the mean, centered on the current value. Whatever is not retained is shared equally across all interactions, including the self term.

Evolutionary simulation results are similar to those from the fixed-sharing model (Supplemental Figure 1). Saturating specialization (i.e., specialization despite concave return function) still occurs for the two-neighbor and bipartite topologies. Thus, for both fixed and evolved resource sharing, we observe specialization for the largest range of parameters (including *α* < 1) not when the group is maximally connected, but rather when connections are fairly sparse. Therefore, a sparse group topology readily constitutes a cooperation-prone physical substrate that can sustain evolvability of specialization traits.

As an example of the benefit of evolving sharing, consider that the maximum fitness according to eq. 1 for a group of *N* disconnected individuals scales as 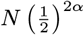. On the other hand, for a bipartite network with a complete specialization strategy (i.e. 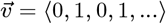), the fitness scales as 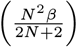. The ratio of these fitnesses is 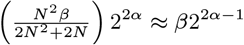, where the approximation is for large *N*. So for larger groups and when 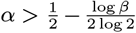, if a group can evolve resource sharing (i.e. letting *β* → 1 and adopting the specialist investment strategy) its maximum fitness will increase.

### Benefit of specialization

We now consider another concrete example to highlight the benefit of specialization. For specialization to be favored, its emergence must result in a higher fitness for the group. To show how this is achieved despite saturating returns, we consider groups of four, connected via the nearest-neighbor topology (i.e., in a ring). We directly calculate the fitness of generalists and specialists for two scenarios: *α* = 0.9 and *α* = 1. By summing the fitness contribution of each member, we can calculate the relative fitness of generalists and specialists, and thus determine whether specialization will be favored (Figure 2).

**Fig. 2.**
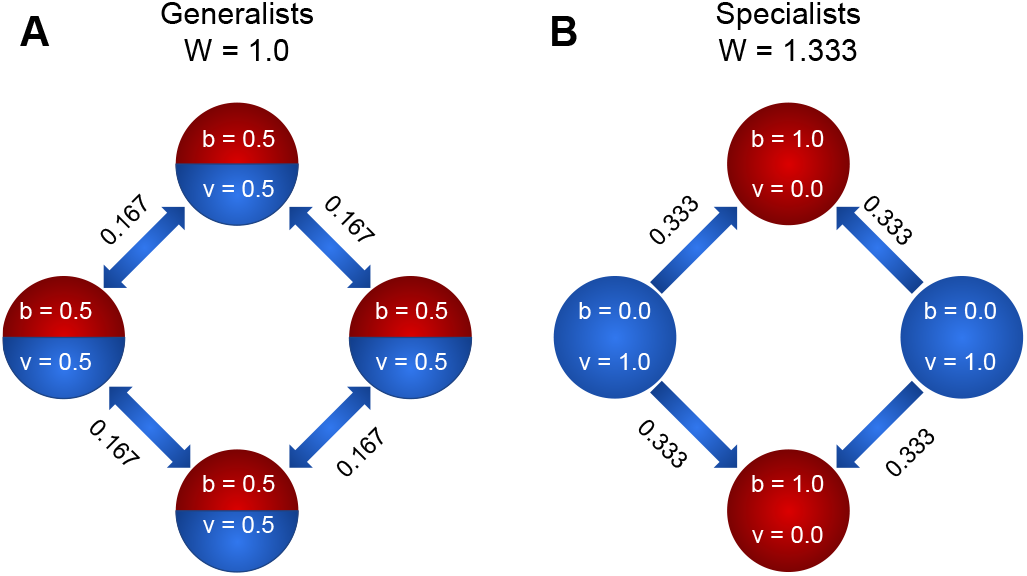
To explore how specialization can be favored by the nearest-neighbor topology, we compare the fitness of a four member system when individuals are (a) generalists and (b) specialists. We first consider the case of linear functional returns (*α* = 1). For the case of generalists (a), each individual receives as much viability as it shares, and all nodes contribute equally to the fitness of the group. Therefore, the fitness of the group is 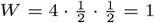. For the case of specialists, however. the viability specialist individuals (blue) have 0 fitness, while the fecundity specialist individuals have nonzero fitness contributions due to the fact that they receive 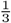 of each viability specialist’s output. Thus the fitness of the group is 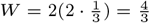. Thus fitness is higher for the group of specialists, so specialization is favored. For *α* = 0.9, the fitness of generalists is 1.15, and the fitness of specialists is 1.39. Thus, even though the returns on investment are saturating (i.e., concave), specialization is favored.

In this simple scenario, reproductive specialization strongly increases group fitness (33% for *α* = 1 and 22% for *α* = 0.9), suggesting an advantage may be found in many systems, even those that are considerably less optimal than the one considered here. For example, real interactions have costs (e.g., transport costs, loss of shared product, etc.); reproductive division of labor would still be favored as long as those costs do not exceed the benefit of specialization.

The benefit of specialization in ring networks increases with group size. For a ring of size *N*, fitness under the specialist strategy 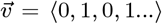 is 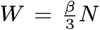. For a ring of generalists the fitness is 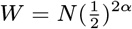. Therefore, whenever 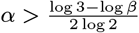, the ring of complete specialists enjoys a greater fitness than the ring of complete generalists. Again, note that complete generalization becomes disfavored when 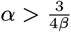, so there *is* a narrow regime where 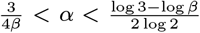 during which neither complete generalization nor complete specialization is optimal. Numerical optimization and evolutionary simulations suggest that even in this region, however, the specialization score of the optimal strategy is large (Figure 1). While these particular topologies do favor specialization even when *α <* 1, we find that the emergence of specialization under these conditions is quite robust to choice of group topology.

### Effect of sparsity

Surprisingly, saturating specialization appears to be the rule, rather than the exception, for sparsely connected graphs. We investigated Erdős-Rényi random graphs with varying degrees of connectivity to systematically examine the relationship between sparsity and the value of *α* at which specialization is favored. We find that many randomly assembled graphs obtain maximum fitness through complete reproductive specialization even when *α* is below 1 (Figure 3 b,c). It is only at the extremes of sparsity and connectivity (near the fully connected or fully unconnected points) that generalists maintain superior fitness for all values of *α <* 1. We further show that this general trend is independent of the size of a group; saturating specialization is favorable for groups of size *N* = 10, *N* = 100, and *N* = 1000. When network connectivity is at its minimum, the group consists solely of isolated individuals that cannot interact. Under these conditions generalists are favored. Similarly, at maximum connectivity every individual interacts with every other individual. Under these conditions generalists are favored unless *αβ* > 1. However, when connectivity is small but not zero, specialization arises most readily. We conjecture that the troughs in Figure 3 c, where specialization occurs for the lowest values of *α*, occur when connectivity is just large enough so that a spanning tree is more likely to connect all individuals in the group than not.

**Fig. 3.**
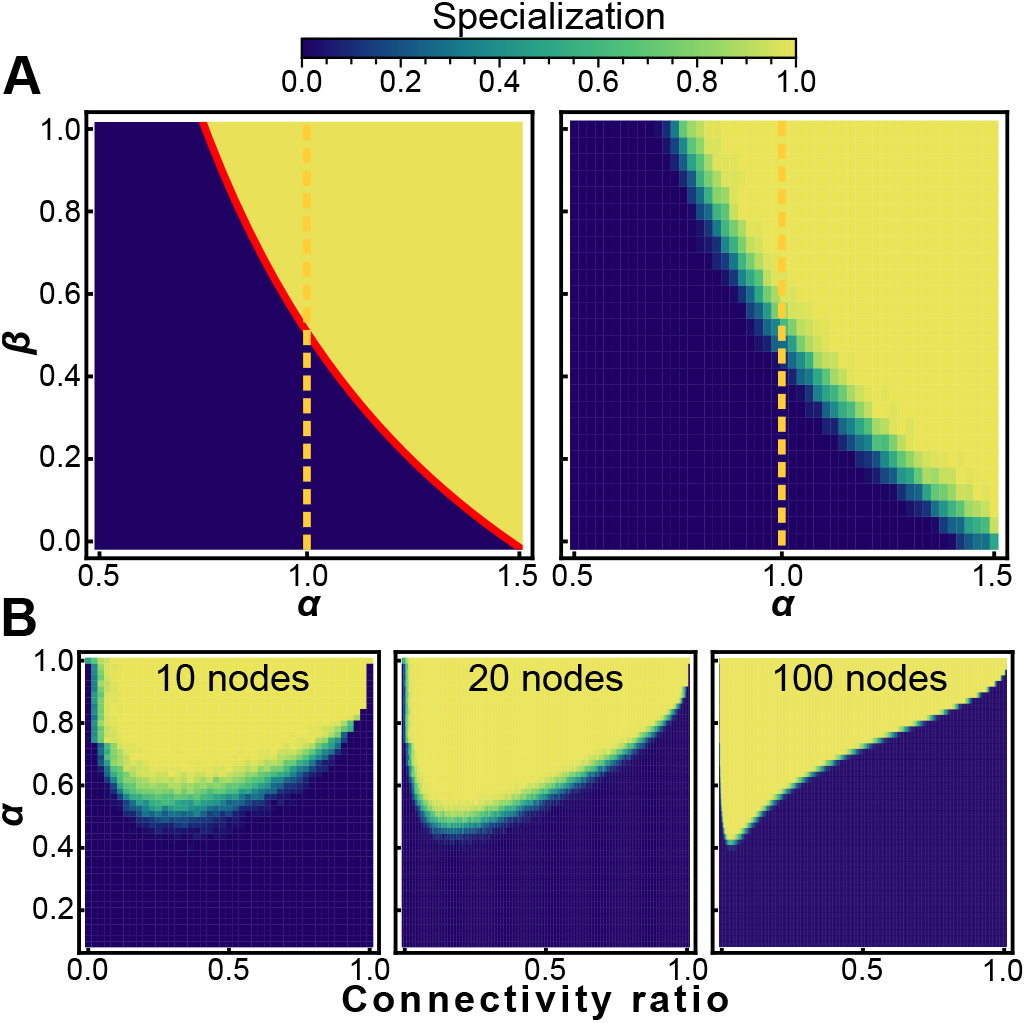
Sparsity encourages specialization. Heat maps showing conditions that favor specialists (white) and generalists (black) for nearest neighbor topologies (A, left) and randomly generated graphs with the same connectivity as nearest neighbor topologies (A, right). Specialization is adaptive on a neighbor network for 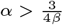; random networks with the same mean connectivity as the nearest neighbor topology behave similarly. (B) The sparsity of a random graph affects how likely it is to favor specialization. We numerically maximize fitness for random graphs of size *N* = 10 (left), *N* = 20 (middle), and *N* = 100 (right) at different levels of sparsity, and subsequently measure the specialization 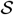 of the fitness maximizing investment strategy. The horizontal axis is the fraction of possible connections present ranging from 0 (none) to 1 (all). The vertical axis is the specialization power *α*, and the colormap shows mean specialization.

### Filaments and trees

Sparse topologies like the two-neighbor configuration have significant biological relevance, and direct ties to early multicellularity. The first step in the evolution of multicellularity is the formation of groups of cells (1, 40–43). Simple groups readily arise through incomplete cell division, forming either simple filaments (Figure 4 a) or tree-like morphologies (Figure 4b) (44–47). Filament topologies have been widely observed in independently-evolved simple multicellular organisms, from ancient fossils of early red algae (48)(Figure 4a) to extant multicellular bacteria (34) and algae (33). Branching multicellular phenotypes have also been observed to readily evolve from baker’s yeast (49)(Figure 4b), and are reminiscent of ancient fungus-like structures (50) and early multicellular fossils of unknown phylogenetic position from the early Ediacaran (45).

**Fig. 4.**
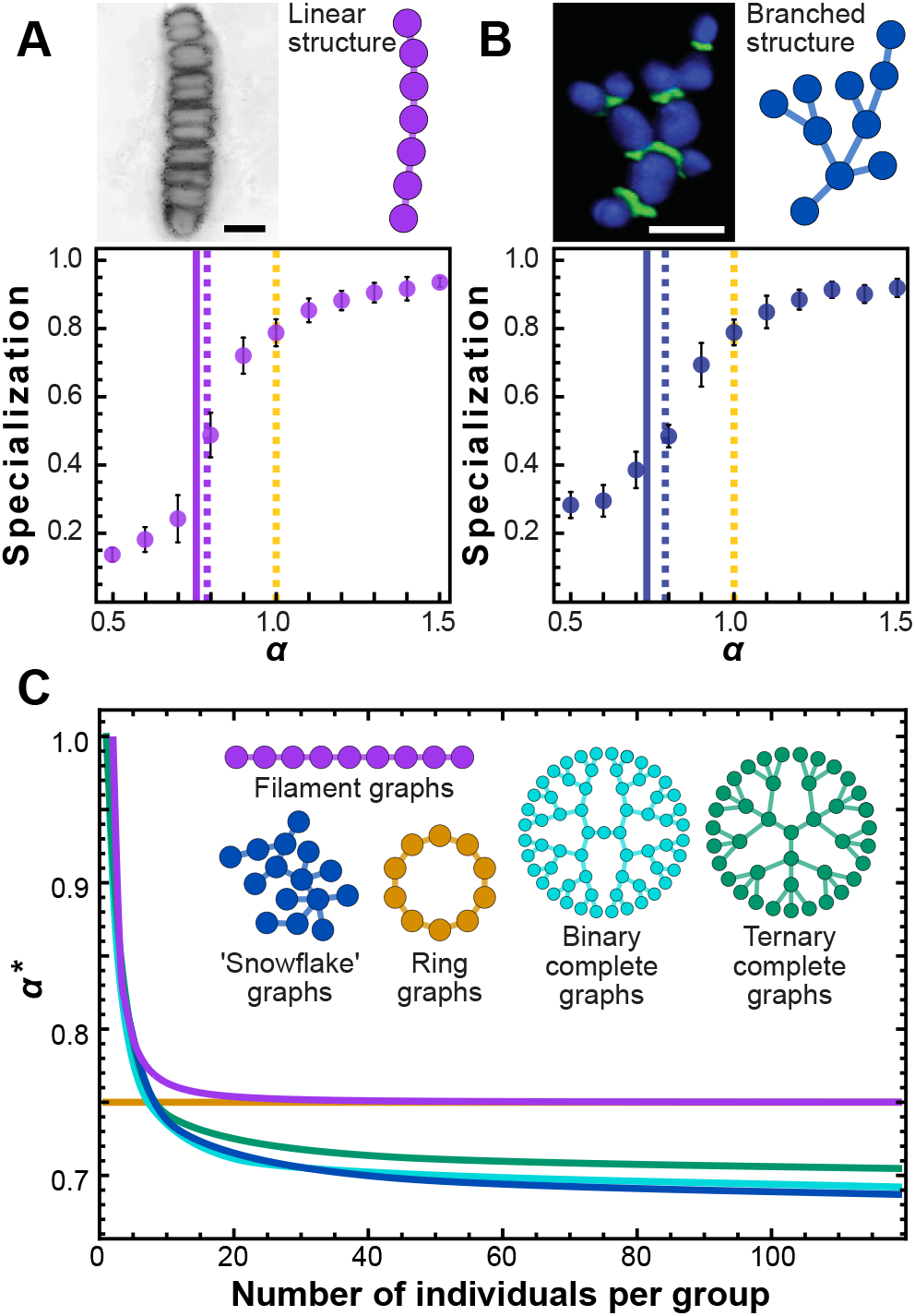
Simple multicellular organisms with sparse topologies. We show two examples of simple multicellular organisms with linear and branched topologies. The image in (a) is a fossilized rhodophyte specimen of *Bangiomorpha pubescens*, courtesy of Prof. Nicholas Butterfield (see e.g. (39); the image in (b) is a confocal image of ‘snowflake yeast’ showing cell volumes in blue and cell-cell connections in green. Scale bars in both panels = 10*μ*m. Panels include cartoons depicting simplified topologies. Topologically similar to the two-neighbor configuration, these configurations yield similar simulation results. Specialization is plotted as a function of *α*. Solid green (a) and blue (b) vertical lines indicate analytical solutions for the transition point where the Hessian evaluated at 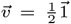 stops being negative definite, i.e. *α*\*; dotted lines indicate roughly where the simulation curves cross specialization of 0.5, i.e. the “true” transition value of *α* where specialization becomes favored. (c) To further explore trees and filaments we analytically solved for *α*\* for various types of trees and filaments of different sizes. *α*\* is plotted versus group size for several topologies. This is a proxy measure of how amenable a network structure is to specialization.

Simulations of populations of groups with filamentous and branched topologies reveal that specialization is indeed favored in the sub-linear regime (Figure 4a and 4b). While the generalist strategy is never a critical point for these networks (which have **c** ≠ **c**^***T***^, see methods), we conjecture that there is a nearby critical point which maximizes fitness at small values of *α* and becomes unstable at larger values of *α*. We introduce a new metric, *α*\*, defined as the value of *α* such that the largest (least negative) eigenvalue of the Hessian evaluated at the complete generalist strategy is zero when *β* = 1. For topologies in which each member has the same number of neighbors, α* is a critical value at which generalization is no longer an optimal strategy. However, even for groups where the number of neighbors for each cell varies, we can still use *α*\* as a proxy for how amenable a topology is to saturating specialization. The smaller *α*\*, the more specialization is likely to be favored. We plot vertical lines where *α* = *α*\* (solid green fig 4(a) and blue fig 4(b)), and dotted lines to indicate roughly where the simulation curves cross specialization of 0.5. This shows that using *α*\* as an overall metric for how amenable a network is to saturating specialization is a reasonable approach— at least in this case. This metric *α*\* only depends on topology and can in principle be calculated analytically given any network. We examined the value of *α*\* as filaments and a variety of tree-like structures grow larger, and find that specialization becomes more strongly favored (Figure 4c). While group size has no effect on specialization for some topologies, like the circular lattice, filaments and trees all see a decrease in *α*\* as group size increases. Once these topologies are larger than a few tens of cells, there appears to be little added benefit to increased group size in terms of specialization. Simple and easily accessible routes to multicellular group formation can readily evolve in response to selection for organismal size (47), and this process may also strongly favor the evolution of cellular differentiation (42, 51–53).

## Discussion

During the evolution of multicellularity, formerly autonomous unicellular organisms evolve into functionally-integrated parts of a new higher-level organism (11, 54). Evolutionary game theory (19, 55, 56) argues that functional specialization should only evolve when increased investment in trade increases reproductive output. Conventionally, this requires returns from specialization to be accelerating, i.e, convex or super-linear (1, 17–25). While this idea is intuitive, it is, in the case of fixed group topology, also overly restrictive. In this paper, we explore how social interactions within groups, measured by their network topology, affect the evolution of reproductive specialization. Indeed, when all individuals within groups interact (with equal interaction strength), benefits must be super-linear for specialization to evolve (figure 1b) (1, 6, 17, 19). Yet for a broad class of sparsely-connected networks, complete specialization can evolve even when the fitness function is saturating, i.e., concave (figure 3).

Rather than being unusual, networks favoring specialization readily arise as a consequence of physical processes structuring simple cellular groups (27). For example, septin defects during cell division create multicellular groups with simple graph structures (Figure 4 a and b), where cells are connected only to parents and offspring (44, 45, 47, 57). If cells share resources only with physically-attached neighbors, then the physical topology of the group describes its interaction topology, and these networks strongly favor reproductive specialization.

Disentangling the evolutionary underpinnings of ancient events is notoriously difficult. Still, it is worth examining the independent origins of complex multicellularity, which are independent runs of parallel natural experiments in extreme sociality. Complex multicellularity (large multicellular organisms with considerable cellular differentiation) has evolved in at least 5 eukaryotic lineages, once each in the animals (58), land plants (59), and brown algae (60), two or three times in the red algae (61, 62), and 8-11 times in fungi (63). In all cases other than animals, these organisms form multicellular bodies via permanent cell-cell bonds, creating long-lasting highly structured cellular networks. Both fossil and phylogenetic evidence suggests that early multicellular organisms in these lineages were considerably less complex, growing as relatively simple graph structures. For example, 1.2 billion year old red algae formed linear filaments of cells (48), basal multicellular charophyte algae formed circular sheets of cells radiating from a common center (59), the ancestor of the brown algae likely formed a branched haplostichous thallus that was either filamentous or pseudoparenchymatous (60), and hyphal fungi are primarily composed of linear chains of cells.

The main difference between our work and previous investigations of the effect of group topology on specialization is that we consider the productivity of groups as a whole, not the individuals within them, and we consider situations of highly asymmetric sharing. Our approach is general, and can be applied to other systems of trade and specialization, so long as 1) only the aggregate productivity of the group (and not the individuals within it) is maximized, 2) the productivity of each individual within the group is a multiplicative function of returns on investment into two (or more) tasks, and 3) there is an asymmetry in how products of those investments are shared. While in this work we have focused on reproductive division of labor, a process in which fecundity returns are not shared at all, we show in the supplement that as long as sharing of two goods is sufficiently asymmetric, specialization with saturating returns on investment can still be adaptive (Supplemental Figure 2).

## Conclusion

We explored the evolution of reproductive specialization in multicellular groups with various cellular interaction topologies. Our results demonstrate that group topological structure can play a key role in the evolution of reproductive division of labor. Indeed, within a broad class of sparsely connected networks, specialization is favored even when the returns from cooperation are saturating (i.e., concave); this result is in direct contrast to the prevailing view that accelerating (i.e., convex), returns are required for natural selection to favor increased specialization (6–11). Our results underscore the central importance of life history trade-offs in the origin of multicellularity (7, 10, 64–66), broadening the conditions under which they can drive the evolution of complete reproductive specialization. Our results support the emerging consensus that evolutionary transitions in individuality are not necessarily highly constrained (4, 43, 47, 65, 67–72).

## Methods

### Analysis

The gradient of the fitness with respect to the group investment strategy 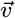, is

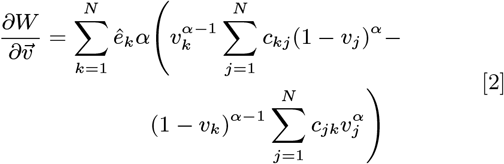

where 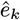 is a unit vector in the *k*^th^ direction. First notice that if **c = c**^***T***^, and 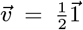 where 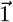 is a vector of ones, then the gradient is zero. This strategy, 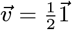, corresponds to the ‘generalist’ approach, where every individual invests equally into both tasks. Call it the generalist strategy. Second, notice that if **c ≠ c**^***T***^then the gradient is *not* zero under the generalist strategy, so at least some degree of specialization must be necessary to maximize fitness. To determine the stability of this solution we examine **H**\*, the Hessian (see SI eq. 6) evaluated at the generalist critical point. If **H**\* is negative definite, then the generalist strategy is a fitness maximum and is therefore an optimal strategy. If, on the other hand, **H**\* has both positive and negative eigenvalues then the generalist strategy lies at a saddle point within the fitness landscape, and therefore the optimal strategy must be somewhere else in (or on the boundary of) the domain (i.e. *v*_*i*_ ∈ [0, 1] for all *i* ∈ 1, 2*, …N*). Finally, note that **H**\* is never positive definite since 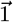 is always an eigenvector with negative eigenvalue (when **c = c**^***T***^).

We also use the zero crossing of the largest eigenvalue of **H**\* evaluated at 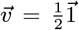 and *β* = 1 as an overall measure of how amenable a network is to specialization, even when **c ≠ c**^***T***^.

### Evolutionary simulations

Our evolutionary simulations maintain the same overall structure as the Wright-Fisher model: a discrete-time Markov chain framework with fitness-weighted multinomial sampling between generations and constant population size. Therefore we refer to them as Wright-Fisher evolutionary simulations. We initialize a population of 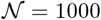 groups, each of group size *N* = 10, with uniform random investment strategies. We then let them evolve for 1000 generations, selecting offspring according to the relative fitness of each group (see eq. 1). At each generation there is a 2% chance for a mutation to a given group’s investment strategy 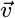. If a mutation occurs, a new investment strategy is selected from a truncated multivariate gaussian distribution centered at the current (pre-mutation) investment strategy and with standard deviation equal to 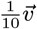. After mutations each group’s fitness is calculated according to eq. 1, and the population is ranked according to fitness. Finally, 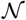 groups are selected (with replacement) to populate the next generation, according to a multinomial distribution weighted by the groups’ fitness ranks.

### Measuring specialization

To quantify the degree of specialization associated with a given group’s optimal investment strategy— the one which maximizes the fitness— we introduce the following metric, which we refer to simply as “Specialization”:

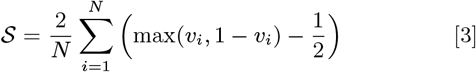

Specialization ranges from 0 (for groups consisting of cells investing equally in functions *v* and *b*) to 1 (for groups consisting of cells investing exclusively in either function

## Code availability

All evolutionary simulations and other computations associated with this work are available at github.com/dyanni3/topologicalConstraintsSpecialization.

## Supplementary Information

### Analysis

As described in the main text, the fitness for a group of *N* individuals is defined as

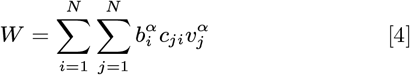

and the gradient of the fitness with respect to the group investment strategy 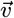, is

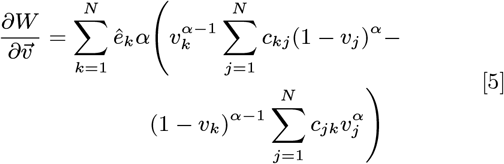

where 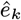 is a unit vector in the *k*^th^ direction.

### Hessian

The Hessian 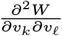 is 

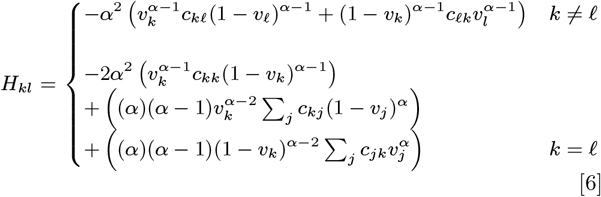

Of particular interest for us is the value of the Hessian at the generalist strategy when **c = c**^***T***^. In that case

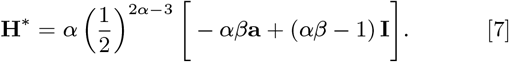

where **a** is the row-normalized adjacency matrix of the network. If **A** is the network’s adjacency matrix then

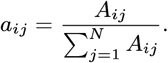

### The case when c = c^*T*^

As noted above, when **c = c**^***T***^, the generalist strategy is always a critical point where 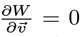. To determine the stability of this solution we examine **H**\* (eq. 7). If **H**\* is negative definite, then the generalist strategy is a fitness maximum and is therefore an optimal strategy. If, on the other hand, **H**\* has both positive and negative eigenvalues then the generalist strategy lies at a saddle point within the fitness landscape, and therefore the optimal strategy must be somewhere else in (or on the boundary of) the domain (i.e. *v*_*i*_ ∈ [0, 1] for all *i* ∈ 1, 2*, …N*). Finally, note that **H**\* is never positive definite (when **c = c**^*T*^). Consider 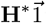:

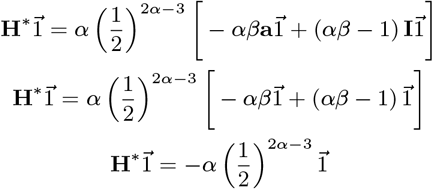

We have 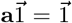 since **a** is row-normalized. Furthermore, *α >* 0, so 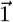 is always an eigenvector of **H**\* with a negative eigenvalue.

We can next ask, under what conditions is **H**\* negative definite? This will depend on the group topology, the nonlinear returns on investment *α*, and the interaction strength *β*. We examine three cases: the ring graph, the bipartite graph, and the complete graph.

When **c = c**^***T***^, the matrix **H**\* is a special type of matrix called a circulant matrix, with well known properties. Its eigenvalues are given by the discrete Fourier transform of its first row. The *k*^th^ eigenvalue is

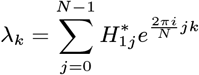

For the ring topology with *N* = 10, for example

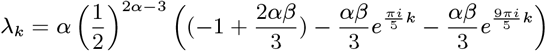

Which has its maximum when *k* = 5,

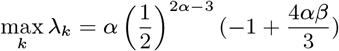

The maximum eigenvalue for the bipartite and complete graphs can be computed similarly.

### Evolution of resource sharing

Here we model the co-evolution of sharing and specialization. We start with generalists that do not share at all. We then allow the amount of sharing and the degree of specialization to evolve. As described in the main text, during every round, each group in the population has a 2% chance that one if its cells will mutate and change how much ‘viability’ it shares. When this occurs, the fraction of its output to retain is chosen from a Gaussian with standard deviation of 10% centered on the current value. Whatever is not retained is shared equally across its interactions. The degree of specialization evolves as in simulations described in the main text.

Results are shown in Figure 1, for nearest neighbor topologies, specialization optimized topologies, and for a complete network.

### General case of sharing two resources

We have so far focused on reproductive specialization, wherein the returns from one type of task (reproduction) are completely unshared while returns from another task (viability) are shared according to some functional interaction strength *β*. Here, we generalize somewhat to consider the returns from two arbitrary tasks which may each be shared to some extent, given by functional interaction strengths (*β*_1_, *β*_2_). For notational continuity we will continue to refer to the investment in those tasks as *b* and *v*, and for tractability we will continue to assume that *α*_2_ = *α*_2_ and that there is a single topology governing who can trade with whom within the group. Of course, further generalizations could be made — e.g. each task could experience different returns on investment, there could be an arbitrary number of tasks, the availability of trading partners could differ between tasks, etc. However, we hope to show by this relatively modest generalization that there is nothing unique to *reproductive* tasks whose fruits are totally unshared that leads to specialization under regimes of sublinear return on investment.

The fitness function is modified so that

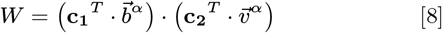

which yields the following Hessian at the generalist critical point (for the ring, bipartite, and complete networks).

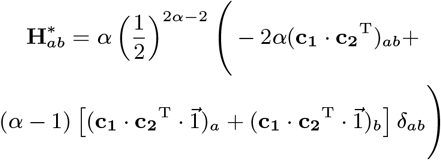

Where

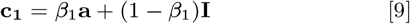

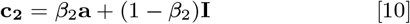

and,

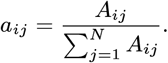

Where, as above, **A** is the graph’s adjacency matrix (including self loops).

**Supplemental Figure 1.**
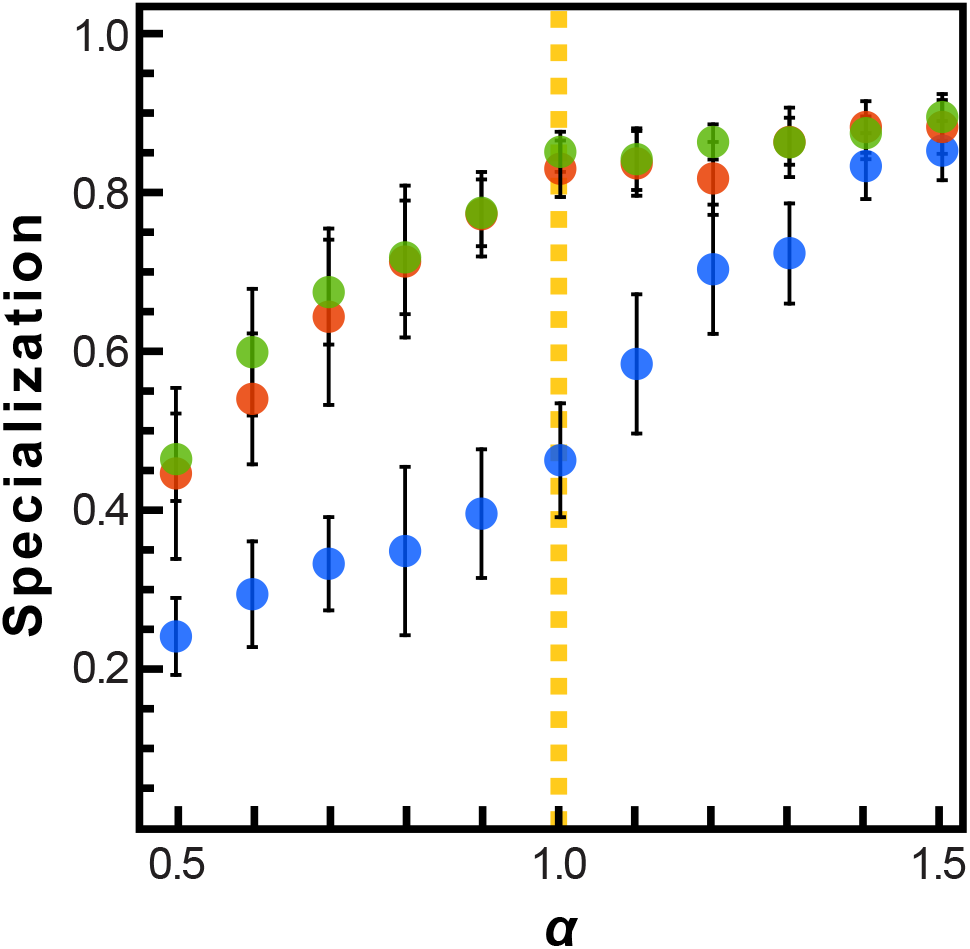
EVOLUTION OF RESOURCE SHARING. Initially, individuals do not share resources; however, they may evolve to do so via random mutations. Here, the mean specialization of the fittest of 100 10-individual groups after 100,000 steps is plotted function of specialization power. Shading is standard deviation across 10 replicates. Blue is the complete network, red is the nearest-neighbor network, and green is the specialization-optimized topology.

We see that for a given topology the adjacency matrix is fixed, so that **c**_**1**_ and **c**_**2**_ differ only in their functional interaction strengths *β*_1_ and *β*_2_. Therefore the maximum fitness strategy, specified by the vector 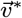, for a given group will depend under our model on the following parameters:

**A** → Adjacency matrix, specifies topology
*β*_1_ → Functional interaction strength of resource 1
*β*_2_ → Functional interaction strength of resource 2
*α* → Specialization power, assumed to be equal for resource 1 and 2

**Supplemental Figure 2.**
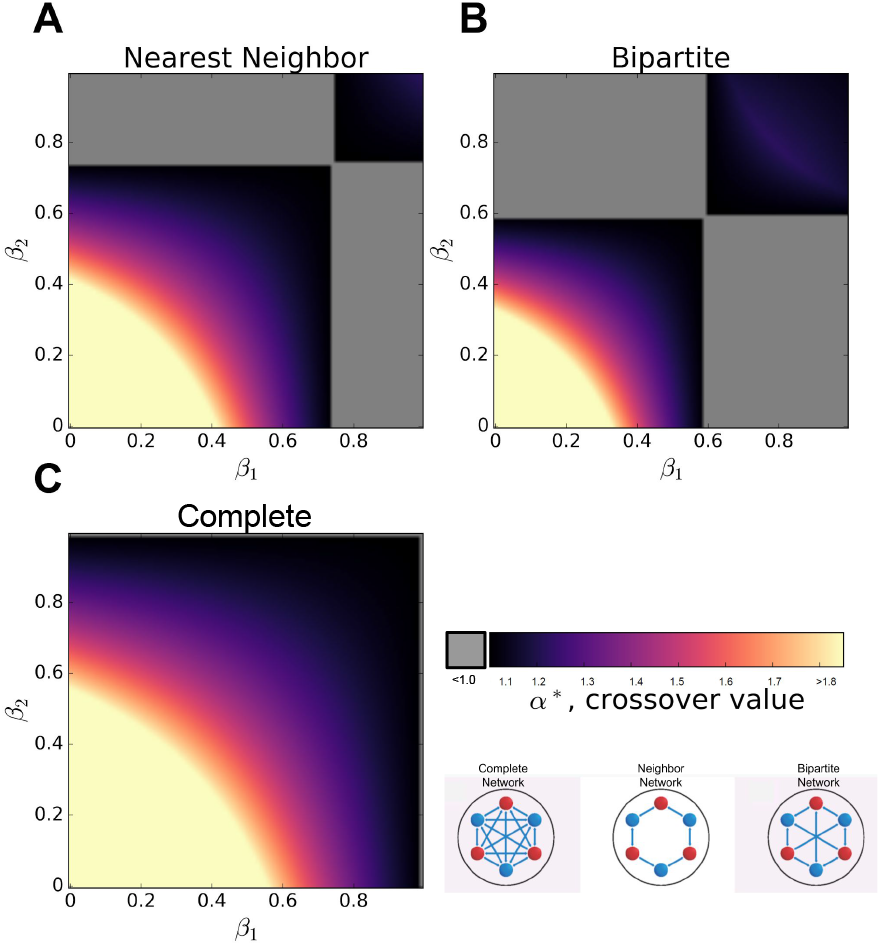
EFFECT OF SHARING TWO RESOURCES. When two resources are shared to different degrees, specified by (*β*_1_, *β*_2_), specialization is sometimes favored under conditions of sublinear returns on investment *α*\* < 1.0. Interestingly, specialization is favored when one resource is shared liberally while the other resource is shared sparingly (though it is not necessary to have one resource remain totally unshared).

We demonstrate the effect of these parameters on the optimal strategy by finding the minimum value of *α* for which specialization becomes favored, which we denote the crossover *α*\*, for a given pair (*β*_1_, *β*_2_) and given topology. The results are shown in figure 2.

